# QTL-Seq Identifies QTL for Relative Electrical Conductivity Associated with Heat Tolerance in Bottle Gourd (*Lagenaria Siceraria*)

**DOI:** 10.1101/2019.12.27.889261

**Authors:** Hui Song, Yunping Huang, Binquan Gu

## Abstract

Heat is a major abiotic stressor seriously affects watermelon produce; however, its effects may be mitigated through grafting onto heat tolerant bottle gourd rootstock. To understand the genetic basis and select reliable markers are necessary in the variety breeding process. In this paper, the relative electric conductivity (REC) was used as a visual indicator for heat stress to study the genetics and SNP marker of heat tolerance in bottle gourd via QTL-seq approach. The results showed that recessive inheritance and a major QTL locus controlled REC related to heat tolerance. Seven heat-tolerant loci (*qHT1.1, qHT2.1, qHT2.2, qHT5.1, qHT6.1, qHT7.1*, and *qHT8.1*) exhibited high Δ(SNP-index) values (ranging from 0.19-0.38), and of these, the greatest value (0.32, P < 0.01) and greatest number of detected SNPs (9052) were found in chromosome 2 at *qHT2.1* region (11.03 - 19.25 Mb); *qHT2.1*was taken as a promising major QTL for heat tolerance in bottle gourd. A total of 34 putative candidate genes related to heat stress were detected within the *qHT2.1* region. Three polymorphic SNPs (BG_GLEAN_10022642, BG_GLEAN_10022727, and BG_GLEAN_10022589) that are involved in pollen sterility, intracellular transport, and signal recognition were validated and exhibited significant marker-trait association that could be used in heat tolerant molecular breeding for bottle gourd. The *qHT2.1*region is an important finding can be used for fine mapping and discovering new genes associated with heat tolerance in bottle gourd.

## Introduction

Heat stress negatively affected physiological processes, reproduction, and adaptation in plants [1], and changes in the climate have exacerbated such effects [2]. Watermelon (*Citrullus lanatus* var. *lanatus*) dominates the fruit market in the world [3], its growth frequently restricted by high temperatures reach more than 35°C during summer months [4]. Bottle gourd (*Lagenaria siceraria* (Mol.) Standl.) tended to be the rootstock for watermelon to reduce heat stress and improve performance [5,6]. Phylogenetic tree analysis has demonstrated the close genetic relatedness of bottle gourd and watermelon [7,8], explains the high grafting survival rate and reduction in taste of bottle gourd in grafted watermelon.

Bottle gourd originated from Africa; it is widely distributed across the tropics [9], where long-term exposure to high temperatures has enhanced its tolerance to heat stress, albeit with wide variation among genotypes [10,11]. Understanding the genetics of heat tolerance in bottle gourd and identification of reliable markers are necessary for the development of novel bottle gourd rootstocks for heat adaptation improvement of scion; however, underlying tolerance characteristics remain unclear.

Heat tolerance is believed to be a polygenic trait that renders the introgression of multiple favorable alleles into cultivars problematic [12]. There have been attempts to deconstruct stress tolerance into measurable components for accurate phenotyping, with the aim that QTLs associated with heat tolerance may then be identified and suitable alleles may be introgressed into elite genetic backgrounds [13]. Cell membrane stability, which has been used as an indicator of heat stress [14], may be quantified by relative electric conductivity (REC). REC is highly sensitive to abiotic stress [15] and has been used in studies of abiotic stress tolerance in a range of crops, including salinity-alkalinity tolerance in muskmelon [16], drought tolerance in *Perennial ryegrass* [17], and cold tolerance in alfalfa [18].

Xu et al.[19] first reported partial sequencing of the bottle gourd genome using the 454 GS-FLX Titanium sequencing platform, from which 400 SSR markers were developed. RAD-Seq [20] technology has also been applied to an F_2_ bottle gourd population for SNP and insertion-deletions marker development [21,22]. Wu et al.[8] reported a high-quality bottle gourd genome sequence was constructed using de novo genome assembly that reconstructed the most recent common Cucurbitaceae ancestor genome through comparison with available extant modern cucurbit species genome resources [23–25].These studies provide an available reference genome and useful molecular markers for the study of bottle gourd agronomic traits.

Many crop plant traits, such as heat stress tolerance, are quantitative, so identification of quantitative trait loci (QTL) may allow efficient crop breeding through marker-assisted selection (MAS) [26]. However, deriving the molecular markers for genetic linkage analysis, QTL analysis is labor-intensive and time-consuming [27]. The bulked-segregant analysis (BSA) provides an effective and simple way to identify molecular markers tightly related to target genes or major QTLs and recent advancements in next-generation sequencing (NGS) technologies provide rapid and efficient QTL identification. For example, Takagi et al. [27] proposed QTL-sequencing (QTL-seq) to detect QTLs that control partial resistance to rice blast using recombinant inbred lines and seedling vigor in F_2_ populations; candidate QTLs detected using the QTL-seq method were confirmed using traditional QTL analysis. The QTL-seq approach has been widely used in a range of crops, such as sunflower [28], rice [29], sorghum [30], potato [31], and cucumber [32,33], for the efficient detection of QTLs for complex quantitative traits. The QTL detection for heat tolerance has been studied in rice [34], chickpea [35],wheat [36], and tomato [37]. In this study, we used QTL-seq to identify the major QTL for heat tolerance in bottle gourd, based on REC, from which polymorphic SNPs and putative candidate genes for heat tolerance were identified using available bottle gourd genome sequence information [21].

## Materials and Methods

### Plant Material and Phenotyping

We selected the heat tolerant inbred line L1 (P17) and sensitive inbred line L6 (P23) as parent material, based on previous work [38], from which F_1_ and F_2_ segregating populations were created. We germinated three replicates of seed from147, 56 and 60 F_2_ individuals of L1, L6 and F_1_ plants, respectively. The seedlings were planted in 32-hole plastic plugs (230cm^3^/hole) filled with nursery substrate (2:1 mix of turf: vermiculite) and grown in a phytotron growth chamber set to a 16 h light/8 h dark cycle (30,000lux) of 25/18°C, with 80% relative humidity. The first true leaf of the seedlings was collected for DNA extraction, and when the third true leaf began to expand, the plants were moved into a high temperature controlled phytotron (40°C, with 80% relative humidity and 30,000lux) for 6h continuous heat exposure.

The electric conductivity (EC) of leaves was measured before and after the heat treatment to assess cell membrane damage, as described by Zhou and Leul [39] and He et al. [40] with modifications. In brief, sampled leaves were washed using deionized water, cut into 0.5-cm pieces, and immersed in deionized water for 30min; then, conductivity of the solution was measured using a conductivity meter (PHSJ-3F, Jingmi Instruments Co., Ltd., Shanghai, China) and recorded as S1. After boiling the leaf samples for 15 min, EC of the solution at room temperature was measured again and recorded as S2. REC was calculated as S1/ S2×100%.

Leaf relative injury (LRI) was used as a metric of cell membrane damage, where LRI = [(Lt-Lck)/(100-Lck)] ×100% [41–43]; Lt and Lck are the REC values before and after heat exposure, respectively, and greater LRI values reflect a lower degrees of heat tolerance.

We used analysis of variance (ANOVA) and t-tests to determine differences in heat tolerance, as indicated by mean (±SE) LRI, among P_1_, P_2_ and F_1_ plants at *P*<0.05 using SPSS 13.0 software [16]. For the F_2_ population, the frequency distribution of LRI was plotted as a histogram and the normal distribution was fitted using the GaussAmp function in ORIGIN 9.1 software [44].

### DNA Sequencing and Analysis

Genome DNA was isolated from young leaves of parental lines L1 and L6, and 147 F_2_ plants using the CATB method [45]. For QTL-seq, two DNA pools (heat tolerance: T-pool; heat sensitive: S-pool) were constructed by mixing an equal amount of DNA from nine heat tolerant (LRI = 1-5%) and nine heat sensitive (LRI = 55-77%) individuals from the F_2_ population.

A library of around 500 bp insert size was constructed and pair-end sequenced (parent average 15×coverage; progeny average was 20×coverage; read length: 150 bp) on an Illumina Hi-Seq4000 at Shanghai Biozeron Co., Ltd.

Raw sequencing data were generated by Illumina base calling software CASAVA v1.8.2 [46] and raw paired end reads were trimmed and quality controlled by Trimmomatic (http://www.usadellab.org/cms/index.php?page=trimmomatic), with default parameters. The high-quality short reads from T-pool and S-pool were aligned to the bottle gourd (Hangzhou gourd) reference genome (http://cucurbitgenomics.org/organism/13, [21]) using BWA software (http://bio-bwa.sourceforge.net/, [47]). After removing PCR-duplicate reads using Picard Tools (http://picard.sourceforge.net/), the valid BAM file was used to detect SNPs using the GATK “UnifiedGenotyper” function (http://www.broadinstitute.org/gatk/); then, low-quality SNPs, with quality value <20 and read depth coverage <4× or >200×, were excluded using custom perl scripts [48].

To identify candidate regions for heat tolerance QTLs, we calculated and plotted two parameters (SNP-index and Δ(SNP-index) for T-pools and S-pools [32,49]. For the polymorphic SNPs located in the genomic regions harboring the major heat tolerance QTL, further functional annotation was completed using available bottle gourd genome data (http://cucurbitgenomics.org/organism/13) and Uniprot database (www.uniprot.org) using ANNOVAR analysis (http://www.openbioinformatics.org/annovar/) to detect putative candidate genes [21].

### SNP Marker Development and Selection

Five candidate SNPs were selected, based on the annotation and classification of putative candidate genes located in the QTL of *qHT2.1* interval; the flank sequences were used for PCR primer development using Primer 6.0 soft (http://www.PromerBiosoft.com). The five SNPs were genotyped for four heat tolerant and four sensitive F_2_ individuals using PCR and Sanger sequencing, and sequence results were visualized and checked using SnapGene software. We used trait-marker association to select the promising SNP markers.

## Results

### LRI Indicator of Heat Tolerance

There were differences in mean LRI between L1 (P17: 9.88±5.00%) and L6 (P23: 47.26±10.78%) under heat stress conditions (Table 1). Mean LRI of the F_1_ was greater (31.98±6.64%) than the mid-parent value (28.32±7.89%), and there was a difference in LRI between F_1_ and L1; there was no difference in LRI between F_1_ and L6. The LRI of the F_2_ progeny ranged from 0.34 to 97.49% (mean: 29.68±8.52) (Table 1). Transgressive segregation was observed on the both sides of the distribution, because distribution of the three F_2_ populations was slightly skewed towards L1 (P17) and showed Gaussian segregation (Fig 1A, B, C).

**Table 1.**
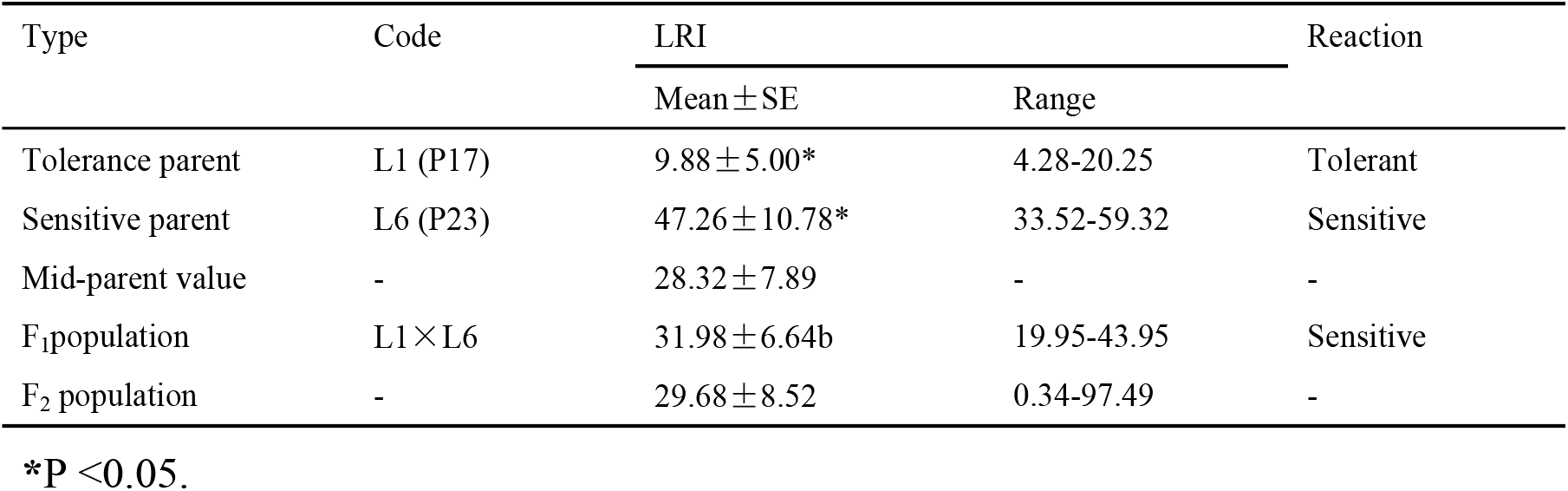
The leaf relative injury (LRI) of parental lines, F_1_ and three F_2_ populations under heat stress conditions.

**Fig 1.**
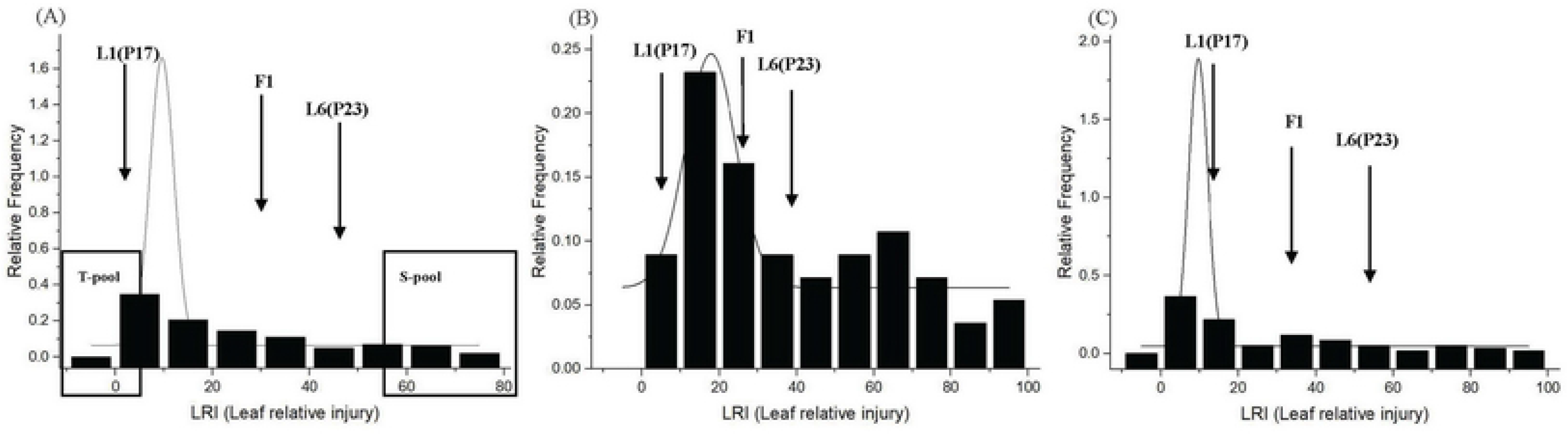
Frequency distribution of leaf relative injury (LRI) shows phenotypic variation among three F_2_ populations with 147 individuals **(A)**, 56 individuals **(B),**and 60 individuals **(C)**. L1: heat tolerant parent P17; L6: heat sensitive parent P23. Distribution of L1 near the origin of the x-axis indicates negative transgressive segregants, while distribution of L6 indicates positive transgressive segregants under heat-stress conditions. DNA of nine F_2_ seedlings was selected from an F_2_ population with 147 individuals **(A)** with extreme phenotypes (low and high LRI values) to develop tolerant and sensitive pools.

### Sequencing Data

Sequencing yielded 31.9, 32.7, 40.9, and 40.9 M 150-bp paired-end raw reads from L1, L6, T-pool, and S-pool, respectively. After trimming and filtering, more than 80% of reads were mapped to the bottle gourd reference genome (Table 2). Aligned to the bottle gourd reference genome (313.4 Mb), we found 23.0, 24.4, 29.0, and 29.5 M short reads were mapped to L1 (10.10×depth coverage or 92.75% coverage), L6 (10.85×depth coverage or 92.85% coverage), T-pool (12.71×depth coverage or 93.25% coverage) and S-pool (12.92×depth coverage or 93.25% coverage), respectively. Based on unique mapped reads, we identified 184,531 and 188,304 SNPs from the T-pool and S-pool, respectively.

**Table 2.** Sequencing and SNP detection.

| Sample | Number of raw reads | Number of trimmed and filtered reads | Uniquely mapped reads |  |  |  | Number of SNPs |  |  |
| --- | --- | --- | --- | --- | --- | --- | --- | --- | --- |
|  |  |  | Number | % | Average Depth | Coverage (%)<sup>a</sup> | Total | Homozygous | Heterozygous |
| L1 | 31,863,484 | 28,368,754 | 22,974,995 | 80.99 | 10.10 | 92.75 | 476,940 | 345,242 | 131,698 |
| L6 | 32,686,118 | 29,026,164 | 24,419,983 | 84.13 | 10.85 | 92.85 | 513,552 | 418,749 | 94,803 |
| T-pool | 40,647,374 | 35,340,578 | 29,026,863 | 82.13 | 12.71 | 93.25 | 543,798 | 359,267 | 184,531 |
| S-pool | 40,930,720 | 35,936,306 | 29,457,284 | 81.97 | 12.92 | 93.25 | 549,415 | 361,111 | 188,304 |

<sup>a</sup>size of the bottle gourd reference genome is 313,397,697 bp (Xu et al. 2014).

### QTL Detection for Heat Tolerance by QTL-seq

The SNP-index graphs were generated for the T-pool (Fig 2A) and S-pool (Fig 2B), and Δ(SNP-index) was plotted against the genome positions (Fig 2C). By examining the Δ(SNP-index) plot, we identified seven genomic positions that exhibited high Δ(SNP-index) values (Table 3) and comprised regions on chromosome 1, from 26.3 to 27.3 Mb (Δ(SNP-index) = 0.25; P < 0.01); chromosome 2, from 11.03 to 19.25 Mb (Δ(SNP-index) = 0.32; P < 0.01) and from 19.4 to 21.0 Mb (Δ(SNP-index) = 0.19; P < 0.01); chromosome 5, from 39.4 to 40.4 Mb (Δ(SNP-index) = 0.38; P< 0.01); chromosome 6, from 7.5 to 8.5 Mb (Δ(SNP-index) = 0.21; P<0.01); chromosome 7, from 23.2 to 24.2 Mb (Δ(SNP-index) = 0.22; P< 0.01); and, chromosome 8, from 24.9 to 25.9 Mb (Δ(SNP-index) = 0.21; P< 0.01). The number of SNPs in the two chromosomes with the greatest Δ(SNP-index) values was greater in Chr2 (11.03-19.25 Mb; 9052) than Chr5 (2) (Table 3) and the SNP-index in the T-pool was lower (0.31) than in the corresponding region of the S-pool (0.63), appearing as a clear mirror image (Δ(SNP-index) = 0.32; P < 0.01; Fig 2). Thus, a promising major QTL controlling heat tolerance appeared to be harbored in the 11.03 to 19.25 Mb regions on chromosome 2 and was designated as *qHT2.1*.

**Fig 2.**
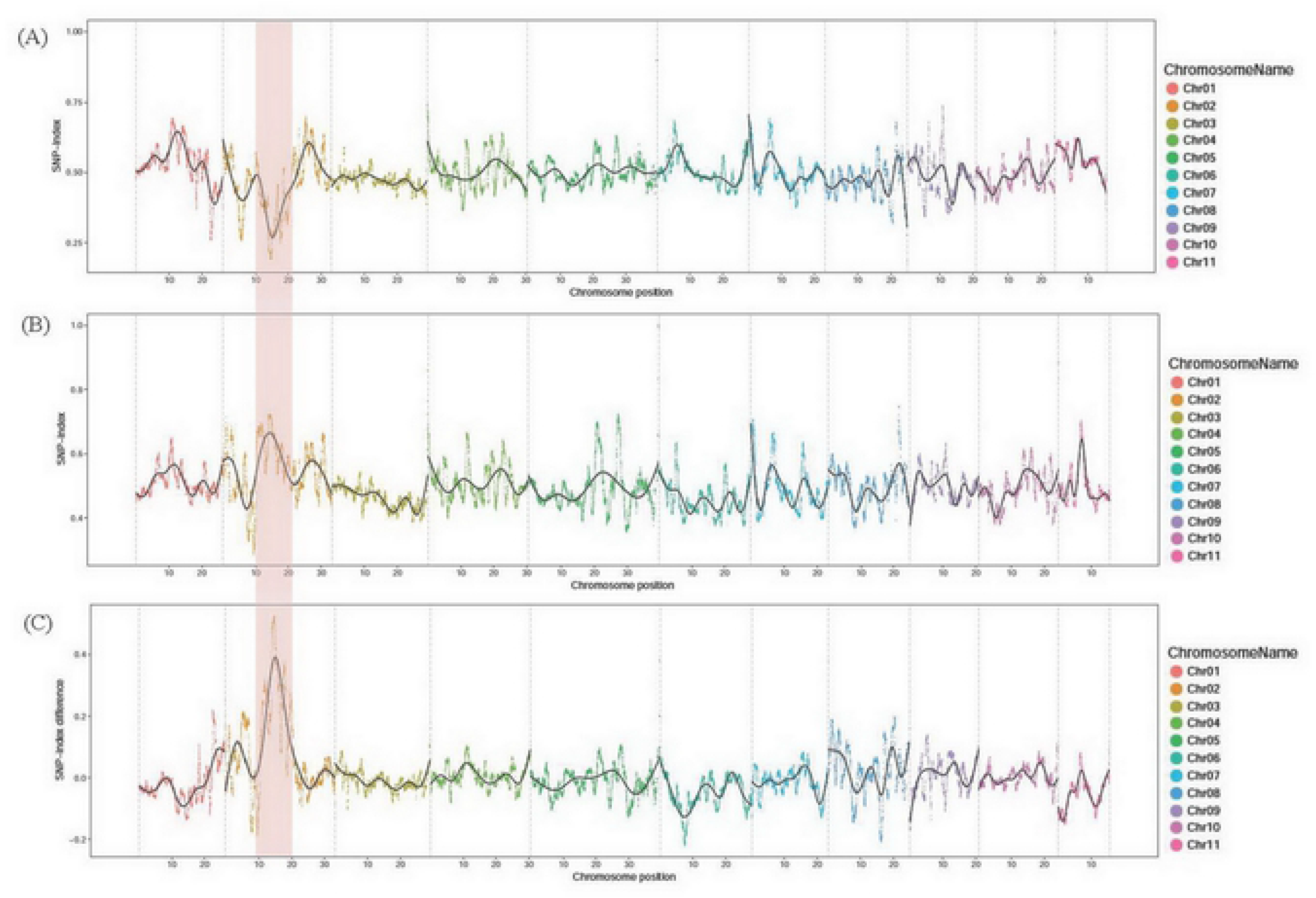
SNP-index plots of T-pool**(A)**and S-pool **(B)** and Δ(SNP-index) plot **(C)** from QTL-seq analysis. The x-axis represents the position of eleven chromosomes and the y-axis represents SNP-index. The Δ(SNP-index) plot (**C**) shows confidence intervals under the null hypothesis of no QTL (P < 0.01). The promising genomic region identified for LRI associated with heat tolerance is shaded at 11.03-19.25 Mb on Chromosome 2.

**Table 3.** QTLs (QTL-seq) that conferred heat tolerance in bottle gourd.

| Region | QTL | SNP<br>Number | SNP-index | | $\Delta$ (SNP-index) | P-value | Interval |
| --- | --- | --- | --- | --- | --- | --- | --- |
|  |  |  | T-pool | S-pool |  |  |  |
| Chr01:26,260,000-27,279,999 | qHT1.1 | 12 | 0.3713 | 0.622 | 0.25 | 0.000413811 | 1,019,999 |
| Chr02:11,030,000-19,249,999 | qHT2.1 | 9052 | 0.3092 | 0.6285 | 0.32 | 9.79E-06 | 8,219,999 |
| Chr02:19,410,000-20,989,999 | qHT2.2 | 1355 | 0.3579 | 0.5463 | 0.19 | 0.006172464 | 1,579,999 |
| Chr05:39,390,000-40,419,999 | qHT5.1 | 2 | 0.45 | 0.8333 | 0.38 | 1.42E-07 | 1,029,999 |
| Chr06:7,460,000-8,459,999 | qHT6.1 | 934 | 0.5917 | 0.379 | -0.21 | 0.001783751 | 999,999 |
| Chr07:23,220,000-24,229,999 | qHT7.1 | 5 | 0.3648 | 0.5812 | 0.22 | 0.001986976 | 1,009,999 |
| Chr08:24,880,000-25,889,999 | qHT8.1 | 4 | 0.1853 | 0.3988 | 0.21 | 0.002248176 | 1,009,999 |

### Identification of Heat Tolerance Candidate Genes in *qHT2.1* Intervals

The full intervals of the regions covered by the seven major Δ(SNP-index) peaks were wide and ranged 8.22 Mb for *qHT*2.1 (Table 3). The 9052 SNPs in this region comprised 527 genes, of which 279 were annotated by BLAST analysis against the non-redundant protein database (S1 Table, [50]) to leave 62 nonsynonymous and stoploss type of SNPs with Δ(SNP-index) ≥0.5 (S2 Table). Gene ontology classification revealed that these 62 genes were mainly associated with biological processes (Fig 3), such as metabolism, cellular components, such as the cell membrane, and molecular function, such as catalytic activity. The biological processes of the 62 candidate genes were retrieved from the available bottle gourd genome data and Uniprot database, 34 genes related to heath stress were determined, along with their corresponding biological processes, comprising pollen and flower sterility, oxidative stress response, autophagy, and abiotic stress response (Table 4).

**Fig 3.**
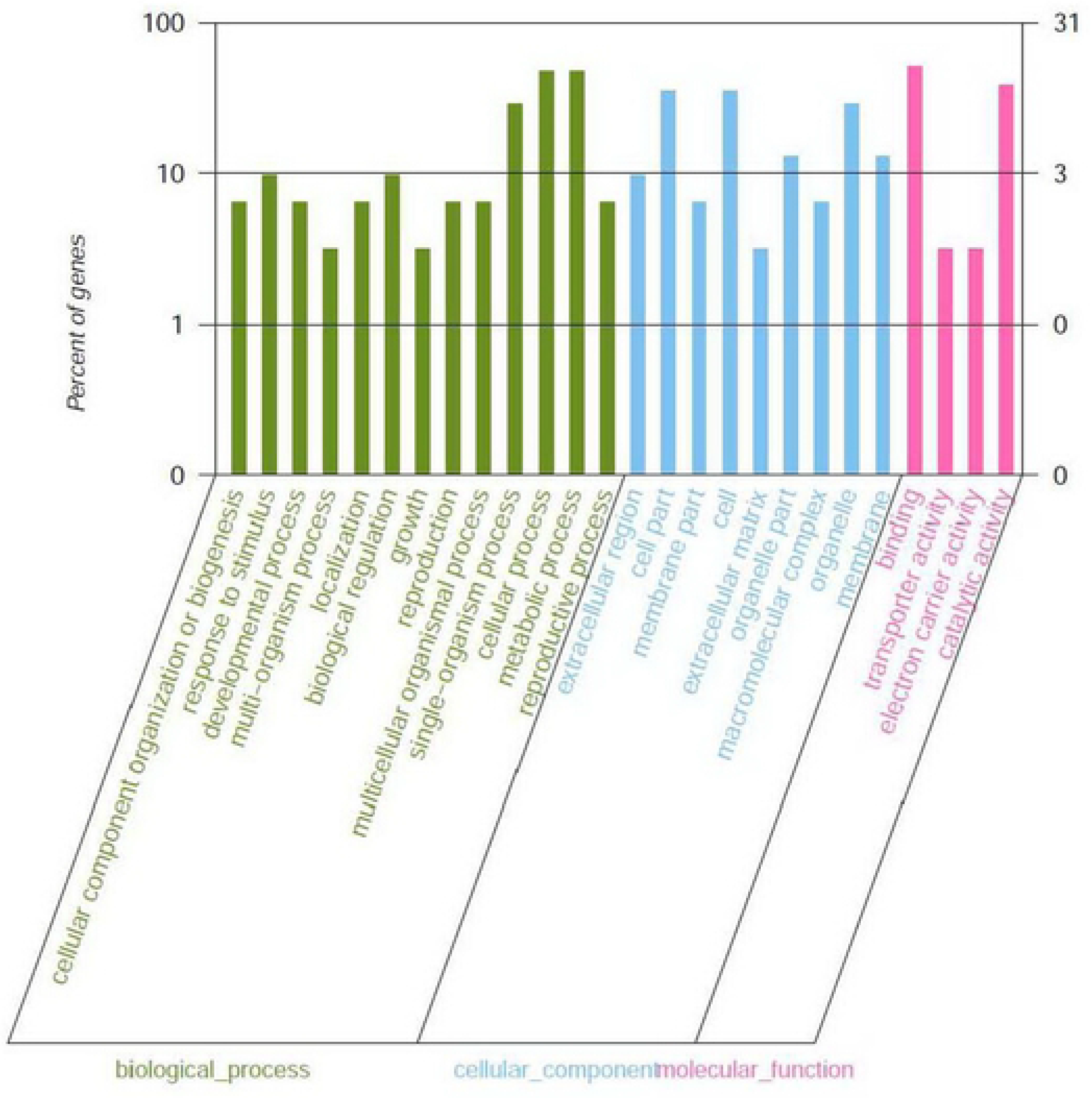
GO results for 62 candidate genes located in the *qHT2.1* region.

**Table 4.**
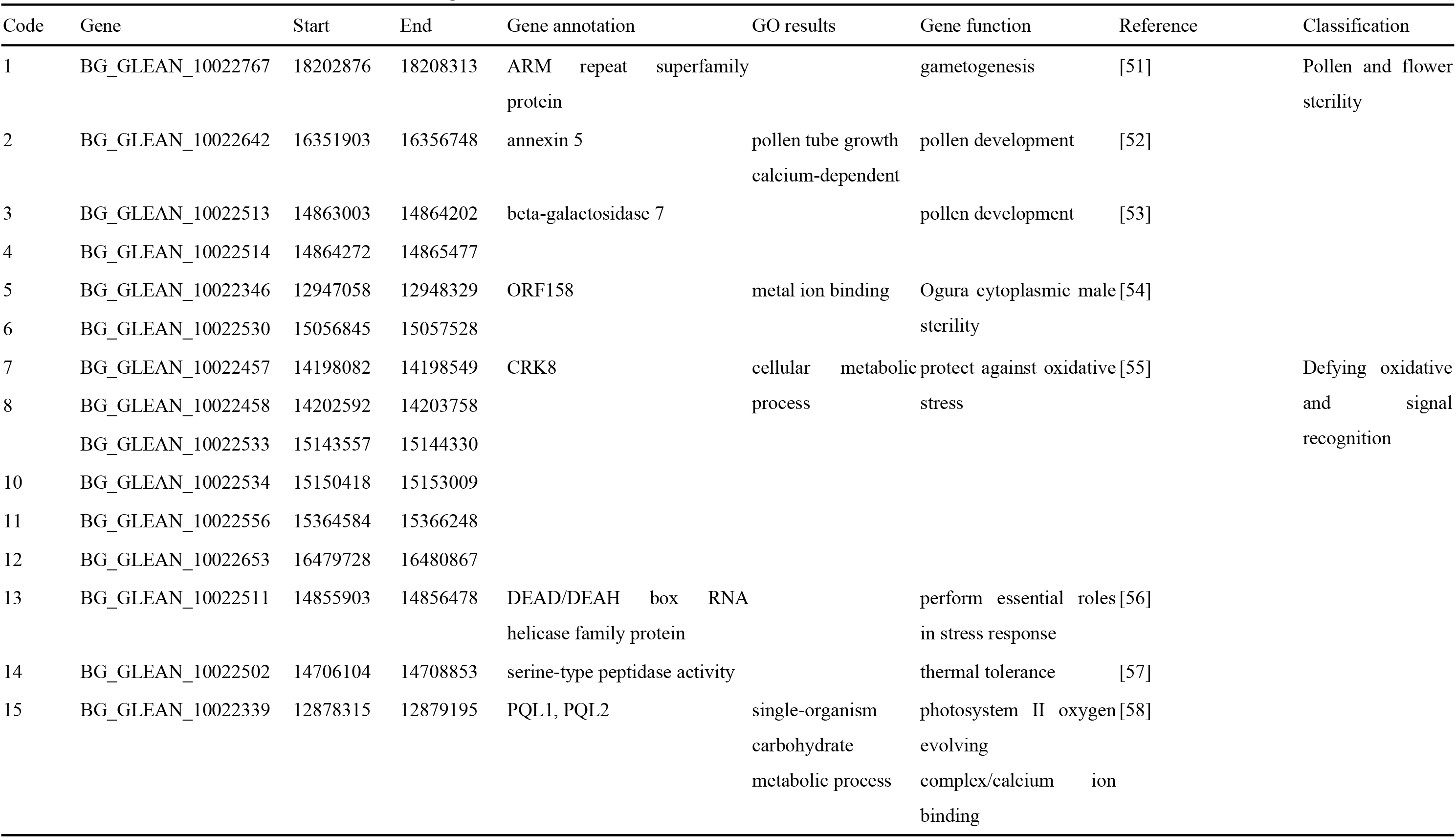
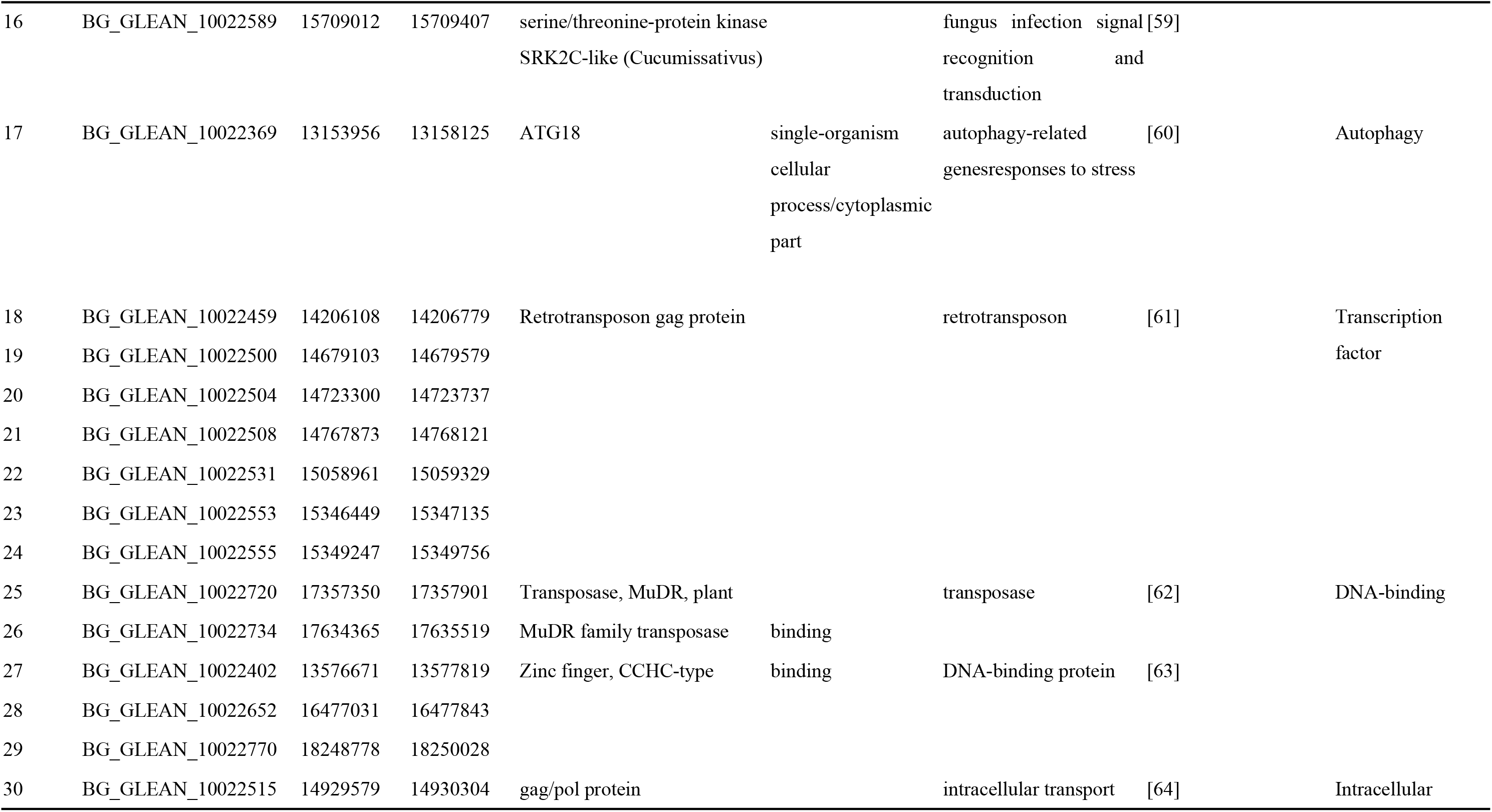
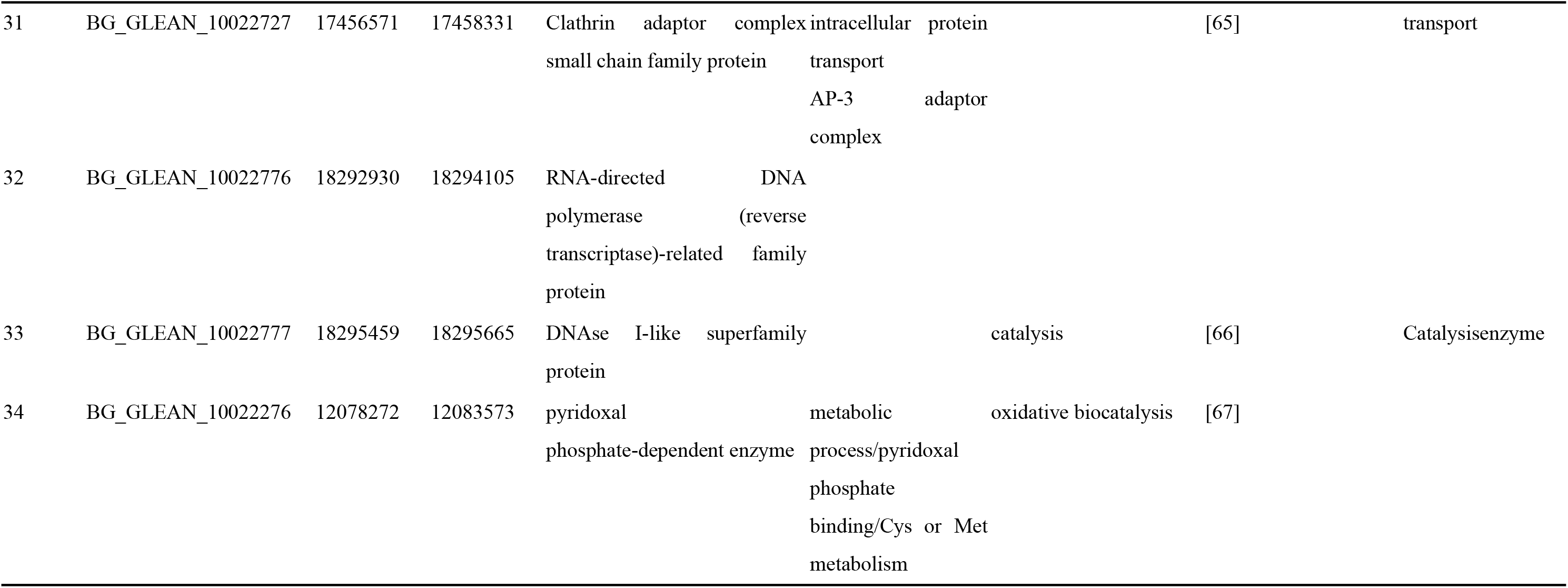
Function annotation of candidate genes.

### SNP Marker Development and Selection

Based on the 34 candidate genes, five SNPs were selected for primer design (Table 5), comprising one related to pollen and flower sterility (SNP 2: BG_GLEAN_10022642, annexin 5), two related to signal recognition (SNP 15: BG_GLEAN_10022339, PQL1 and PQL2; and, SNP 16: BG_GLEAN_10022589, serine/threonine-protein kinase), and two related to intracellular transport (SNP 26: BG_GLEAN_10022734, MuDR; and, SNP 31: BG_GLEAN_10022727, AP-3 adaptor complex). Genotyping of four heat tolerant and four sensitive F_2_ individuals (Fig 4, Table 5) showed two of the five SNPs (SNP 15 and SNP 26) were similar between the two heat sensitivity groups, while SNPs 2 and 31were homozygous in heat tolerant individuals (CC and TT, respectively) and homozygous and heterozygous in heat sensitive individuals (AA and AC for SNP 2, and AA and AT for SNP 31). The only homozygous SNP for heat tolerance and heat sensitive individuals was SNP16 (BG_GLEAN_10022589; GG and AA, respectively. Three polymorphic markers (SNPs 2, 31, and 16) showed good marker-trait association, where SNPs 2 and 31may be dominant markers, and SNP 16 may be a co-dominant marker for heat tolerance.

**Table 5.**
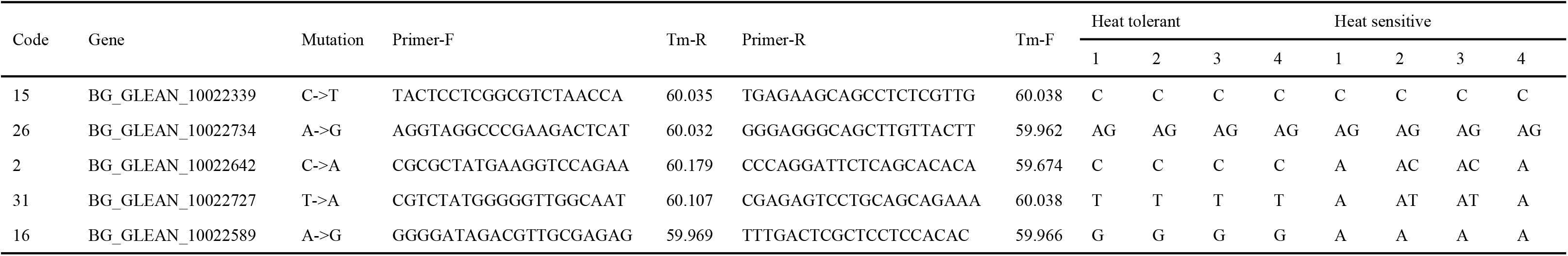
Primer design and Sanger sequencing verification of five candidate loci in four tolerant and sensitive F_2_ individuals.

**Fig 4.**
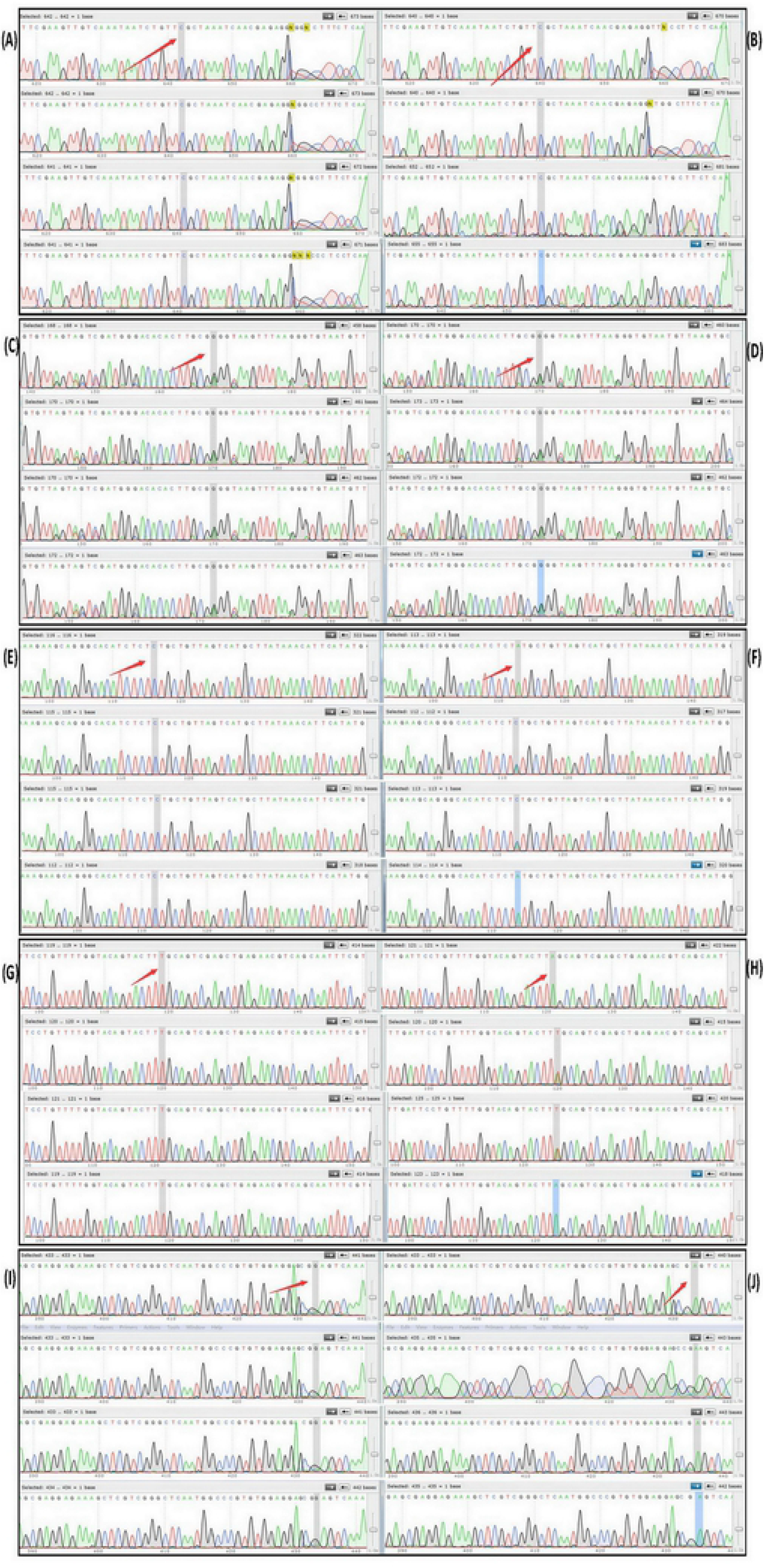
Genotyping of four heat F_2_ tolerant (**A**, **C**, **E**, **G**, **I**) and four heat sensitive F_2_ (**B**, **D**, **F**, **H**, **J**) individuals derived from five candidate SNPs. **(AB)** SNP15: BG_GLEAN_10022339. **(CD)** SNP26: BG_GLEAN_10022734. **(EF)** SNP 2: BG_GLEAN_10022642. **(GH)** SNP 31: BG_GLEAN_10022727. **(IJ)** SNP 16: BG_GLEAN_10022589. The SNP loci are shaded and indicated with a red arrow.

## Discussion

### Genetics of Heat Tolerance in Bottle Gourd

Plants have evolved a range of metabolic responses, such as antioxidant activity, membrane lipid unsaturation, protein stability, gene expression and translation, and accumulation of compatible solutes [68], to cope with heat stress through the activation of stress-response genes [69]. Heat tolerance, which is characterized by a series of phenotypic and physiological traits, may be summarized as the quantitative trait [70]. Poly genes are known to play a critical role in the control of heat tolerance in crop plants, such as wheat [71,72], rice [34], and broccoli [73,74], although there are reports monogenic and oligogenic responses to heat tolerance [75,76]. Previous understanding of inheritance patterns of heat tolerance in bottle gourd was limited, and our study is the first to report on the recessive inheritance and a major QTL locus for heat tolerance in this crop plant, based on REC using a QTL-seq approach.

The plants in the F_1_ progeny of the L1×L6 cross were heat sensitive, indicating that heat tolerance is a recessive trait in bottle gourd, as confirmed through genotyping of the candidate marker SNPs 2 (BG_GLEAN_10022642) and 31 (BG_GLEAN_10022727) as homozygous CC or TT, respectively, that indicate heat tolerance, and A_ (AA, AC or AT) that indicate heat sensitivity. These results also infer the presence of a dominant locus of AA in heat sensitive parent material.

The distribution of the F_2_ population was skewed towards L1, indicating that a major QTL locus controlled heat tolerance-related LRI. According to the principle of QTL-seq, described by Takagi et al. [27], the SNP-index graphs of the T-pool and S-pool were expected to be identical for the genomic regions, without phenotypic differences (heat tolerance), whereas the genomic region(s) harboring heat tolerance QTL(s) were expected to exhibit unequal contributions from P1 and P2 parental genomes. Furthermore, the SNP-indexes of these regions for T- and S-pools should appear as a mirror image around SNP-index = 0.5. Although seven QTLs were identified using QTL-seq, only *qHT2.1* was identified as the most promising major QTL for heat tolerance, because of the greatest Δ(SNP-index) value (0.32, P <0.01) and number of detected SNPs (9052), and clear mirroring of SNP-index plots. Similar results have been reported for African rice (*Oryza glaberrima* Steud.), in which a major QTL that controlled survival rate of seedlings exposed to high temperatures was governed by the Thermo tolerance 1 gene encoded in the 23S proteasome subunit, which degrades ubiquitinated cytotoxic denatured proteins that are formed due to high temperature stress [77]. Major QTL control of complex traits, such as heat tolerance, may be explained by inheritance patterns of the trait [13] and action of major regulator genes that may switch off subordinate genes, if a key gene is mutated [78].

### Annotation of Heat Tolerance Candidate Genes

Understanding of functional genomics in plants facilitates the elucidation of candidate genes and their relationship with traits [35]. Heat stress affected pollen and flower sterility [79]. Of the 34 candidate genes, four genes were identified to have key roles in gametogenesis [51], pollen development [52,53] or Ogura cytoplasmic male sterility[54]. Heat-stress also caused cellular damage by oxidative stress and toxicity due to the reactive oxygen species (ROS) formation [80,81]. Antioxidants scavenge reactive oxygen species to mitigate oxidative. Four putative candidate genes were found to have a function in defying oxidative stress and recovering plants from heat-stress damage. Many transcription factors, binding factors, intracellular transporters and enzyme responsible for abiotic tolerance in different crops were reported earlier [35]. In addition, signalling molecule [59], autophagy [60] play important roles in plant responses to stress.

### Annotation of Candidate SNP

In this study, three SNPs with contrasting function annotation were found to be associated with heat tolerance in bottle gourd, including SNP2 (BG_GLEAN_10022642; annexin 5) associated with pollen and flower sterility; SNP 31 (BG_GLEAN_10022727; AP-3 adaptor complex) related to intracellular transport; and, SNP 16 (BG_GLEAN_10022589; serine/threonine-protein kinase) involved in signal recognition; which exhibited significant marker-trait association and could be attractive option to identify the heat tolerance bottle gourd genotypes.

A common response of crop plants to temperature stress is significant yield loss, as a result of effects on spikelet fertility in rice [82], pot set in lentil [78], and pollen sterility in canola [79]. Annexins belong to a ubiquitous family of proteins present in eukaryotic organisms [83] localized in various subcellular compartments [84], and are known to be involved in a variety of cellular processes, such as maintenance of vesicular trafficking, cellular redox homeostasis, actin binding, and ion transport [85], due to their calcium- and membrane-binding capacity. Annexin 5 is involved in pollen grain development and germination, and pollen tube growth through the promotion of calcium-modulated endo-membrane trafficking [86]. For example, down-regulation of Arabidopsis annexin5 in transgenic Ann5-RNAi lines has been shown to cause sterility in pollen grains [52].

Adaptor proteins that are involved in protein trafficking and sorting [87–89] may recognize cargo and coat proteins during vesicle formation [90]. AP-3 was first identified in mammalian cells and likely functions as a clathrin adaptor [91]; losses in AP-3 function reduce seed germination potential [92,93], due to mistargeted protein S-ACYL transferase10 that is critical for pollen tube growth during dynamic vacuolar organization in Arabidopsis [94].

Serine/threonine-protein kinases (STKs) are involved in signal transduction networks to coordinate growth and differentiation of cell responses to extracellular stimuli [95]. In a range of smut pathogenesis-related biological processes, Huang et al. [59] reported that STKs may act as receptors or signaling factors, such as in Ca^2+^ signaling, that then activate defense responses. Umezawa et al. [96] reported that SRK2C is annotated as a putative and osmotic-stress-activated STK in Arabidopsis; it may mediate signal transduction and regulate a series of drought stress-response genes that enhance expression of 35S:SRK2C-GFP to improve drought tolerance in plants.

## Conclusion

To understand genetic basis and select reliable markers are important in the variety breeding process for bottle gourd. The accurate QTL-seq results have clearly shown that the recessive heat tolerance inheritance and three polymorphic SNPs involved in pollen sterility, intracellular transport, and signal recognition were exhibited significant marker-trait association. Intensity of global warming will increase in next two to three decades and will exacerbate the negative effects on Cucurbitaceae plants. It will help in identification of heat tolerance genotypes in breeding program of bottle gourd for further heat tolerance strengthening of other Cucurbitaceae plants by grafting, such as those for watermelon. The results also revealed the novel region (from 11.03 to 19.25 Mb on chromosome 2) harbored *qHT2.1*, which first provide the promising direction for further exploration on heat tolerance in bottle gourd for fine mapping to discover the new gene and understanding the complex mechanisms.

## Supporting information

**S1 Table. Predicted and annotated SNPs in the *qHT2.1* region.**

**S2 Table. Details of 62 nonsynonymous and stoploss type of SNPs in the *qHT2.1* region.**

## Acknowledgments

We are grateful to Q. F. Lou and Y. Q. Weng for their critical comments on this work. Funding for this research comes from the National Natural Science Foundation of China (NSFC: 31572142 and 31000907) and Special Fund for Agro-Scientific Research in the Public Interest (201403032).

## Author Contributions

**Conceptualization:** Hui Song

**Data curation:** Hui Song, Yunping Huang

**Formal analysis:** Hui Song, Yunping Huang, Binquan Gu

**Funding acquisition:** Hui Song

**Investigation:** Hui Song, Binquan Gu

**Methodology:** Hui Song, Yunping Huang

**Project administration:** Hui Song

**Resources:** Hui Song

**Software:** Hui Song

**Supervision:** Hui Song

**Validation:** Hui Song, Yunping Huang

**Visualization:** Hui Song, Binquan Gu

**Writing - original draft:** Hui Song

**Writing - review & editing:** Hui Song, Yunping Huang

## References

1. Hall JL. Cellular mechanisms for heavy metal detoxification and tolerance. J Exp Bot. 2002; 53: 1–11.

2. Fedoroff NV, Battisti DS, Beachy RN, Cooper PJ, Fischhoff DA, Hodges CN, et al. Radically rethinking agriculture for the 21st century. Science. 2010; 327: 833–834.

3. Tian SW, Jiang LJ, Cui XX, Zhang J, Guo SG, Li MY, et al. Engineering herbicide-resistant watermelon variety through CRISPR/Cas9-mediated base-editing. Plant Cell Rep. 2018; 37: 1353–1356.

4. Brinen GH. Plant and row spacing, mulch and fertilizer rate effects on watermelon production. J Am Soc Hort Sci. 1979; 104: 724–726.

5. Lee JM, Oda M. Grafting of herbaceous vegetable and ornamental crops. Hortic Rev Am Soc Hortic Sci. 2003; 28: 61–124.

6. Yang Y, Lu X, Yan B, Li B, Sun J, Guo S, et al. Bottle gourd rootstock-grafting affects nitrogen metabolism in NaCl-stressed watermelon leaves and enhances short-term salt tolerance. J Plant Physiol. 2013; 170: 653–661.doi: 10.1016/j.jplph.2012.12.013

7. Schaefer H, Heibl C, Renner SS. Gourds afloat: a dated phylogeny reveals an Asian origin of the gourd family (Cucurbitaceae) and numerous oversea dispersal events. Proc R Soc London B Biol Sci. 2009; 276: 843–851.

8. Wu S, Shamimuzzaman M, Sun HH, Salse J, Sui XL, Wilder A, et al. The bottle gourd genome provides insights into Cucurbitaceae evolution and facilitates mapping of a Papaya ring-spot virus resistance locus. Plant J. 2017; 92: 963–975.

9. Erickson LD, Smith BD, Clarke AC, Sandweiss DH, Tuross N. An Asian origin for a 10,000-year-old domesticated plant in the Americas. Proc Natl Acad Sci USA. 2005; 102: 18315–18320.

10. Samadia DK. Performance of bottle gourd genotypes under hot arid environment. Indian J Hort. 2002; 59: 167–170.

11. Mashilo J, Shimelis H, Odindo A. Yield-based selection indices for drought tolerance evaluation in selected bottle gourd (*Lagenaria siceraria* (Molina) *Standl.*) landraces. Acta Agr Scand S-B P. 2017; 67: 43–50. doi: 10.1080/09064710.2016.1215518

12. Asthir B. Protective mechanisms of heat tolerance in crop plants. J Plant Interact. 2015; 10: 202–210.

13. Gilliham M, Able JA, Roy SJ. Translating knowledge about abiotic stress tolerance to breeding programmes. Plant J. 2017; 90: 898–917.

14. Ibrahim E, Saadalla M, Baenziger S, Bockelman H, Morsy S. Cell membrane stability and association mapping for drought and heat tolerance in a worldwide wheat collection. Sustainability. 2017; 9: 1606–1022. doi:10.3390/su9091606

15. Arvin MJ, Donnelly DJ. Screening potato cultivars and wild species to abiotic stresses using an electrolyte leakage bioassay. J. Agr Sci Tech. 2008; 10: 33–42.

16. Jin XQ, Liu T, Xu JJ, Gao ZX, Hu XH. Exogenous GABA enhances muskmelon tolerance to salinity-alkalinity stress by regulating redox balance and chlorophyll biosynthesis. BMC Plant Bio. 2019; l19: 48–65. doi:10.1186/s12870-019-1660-y

17. Su AY, Niu SQ, Liu YZ, He AL, Zhao Q, Paré PW, et al. Synergistic effects of Bacillus amyloliquefaciens (GB03) and water retaining agent on drought tolerance of *Perennial ryegrass*. Int J Mol Sci. 2017; 18: 2651–2664. doi:10.3390/ijms18122651

18. Sulc RM, Albrecht KA, Duke SH. Leakage of intracellular substances as an indicator of freezing injury in alfalfa. Crop Sci. 1991; 31: 430–435.

19. Xu P, Wu XH, Luo J, Wang BG, Liu YH, Ehlers JD, et al. Partial sequencing of the bottle gourd genome reveals markers useful for phylogenetic analysis and breeding. BMC Genomics. 2011; 12: 467–480.

20. Baird NA, Etter PD, Atwood TS, Currey MC, Shiver AL, Lewis ZA, et al. Rapid SNP discovery and genetic mapping using sequenced RAD markers. PLoS One. 2008; 3: e3376. Doi: 10.1371/journal.pone.0003376

21. Xu P, Xu SZ, Wu XH, Tao Y, Wang BG, Wang S, et al. Population genomic analyses from low-coverage RAD-Seq data: A case study on the non-model cucurbit bottle gourd. Plant J. 2014; 77: 430–442.

22. Wu XY, Xu P, Wu XH, Wang BG, Lu ZF, Li GJ. Development of insertion and deletion markers for bottle gourd based on restriction site-associated DNA sequencing data. Hort Plant J. 2017; 3: 13–16.

23. Huang SW, Li RQ, Zhang ZH, Li L, Gu XF, Fan W, et al. The genome of the cucumber, *Cucumissativus L*. Nat Genet. 2009; 41: 1275–1281.

24. Garcia-Mas J, Benjak A, Sanseverino W, Bourgeois M, Mir G, González VM, et al. The genome of melon (*Cucumis melo* L.). Proc Natl Acad Sci USA. 2012; 109: 11872–11877.

25. Guo SG, Zhang JG, Sun HH, Salse J, Lucas WJ, Zhang HY, et al. The draft genome of watermelon (*Citrullus lanatus*) and resequencing of 20 diverse accessions. Nat Genet. 2013; 45: 51–58.

26. Paterson AH, Lander ES, Hewitt JD, Peterson S, Lincoln SE, Tanksley SD. Resolution of quantitative traits into Mendelian factors by using a complete linkage map of restriction fragment length polymorphisms. Nature. 1988; 335: 721–726.

27. Takagi H, Abe A, Yoshida K, Kosugi S, Natsume S, Mitsuoka C, et al. QTL-seq: rapid mapping of quantitative trait loci in rice by whole genome resequencing of DNA from two bulked populations. Plant J. 2013; 74: 174–183.

28. Livaja M, Wang Y, Wieckhorst S, Haseneyer G, Seidel M, Hahn V, et al. BSTA: a targeted approach combines bulked segregant analysis with next generation sequencing and de novo transcriptome assembly for SNP discovery in sunflower. BMC Genomics. 2013; 14: 628–638.

29. Yang Z, Huang D, Tang W, Zheng Y, Liang K, Cutler AJ, et al. Mapping of quantitative trait loci underlying cold tolerance in rice seedlings via high-throughput sequencing of pooled extremes. PLoS One. 2013; 8: e68433. doi:10.1371/journal.pone.0068433

30. Han Y, Lv P, Hou S, Li S, Ji G, Ma X, et al. Combining next generation sequencing with bulked segregant analysis to fine map a stem moisture locus in sorghum (*Sorghum bicolor* L. *Moench*). PLoS One. 2015; 10: e0127065. doi:10.1371/journal.pone.0127065

31. Kaminski KP, Korup K, Andersen MN, Sonderkaer M, Andersen MS, Kirk HG, et al. Next generation sequencing bulk segregant analysis of potato support that differential flux into the cholesterol and stigmasterol metabolite pools is important for steroidal glycoalkaloid content. Potato Res. 2016; 59: 81–97.

32. Lu H, Lin T, Klein J, Wang S, Qi J, Zhou Q, et al. QTL-seq identifies an early flowering QTL located near flowering locus T in cucumber. Theor Appl Genet. 2014; 127: 1491–1499.

33. Win KT, Vegas J, Zhang CY, Song K, Lee S. QTL mapping for downy mildew resistance in cucumber via bulked segregant analysis using next-generation sequencing and conventional methods. Theor Appl Genet. 2017; 130: 199–211.

34. Shanmugavadivel P S, Amitha MSV, Chandra P, Ramkumar MK, Tiwari R, Mohapatra T, et al. High resolution mapping of QTLs for heat tolerance in rice using a 5K SNP array. Rice. 2017; 10: 28–39. doi: 10.1186/s12284-017-0167-0

35. Paul PJ, Samineni S, Thudi M, Sajja SB, Rathore A, Das RR, et al. Molecular mapping of QTLs for heat tolerance in chickpea. Mol Sci. 2018; 19: 2166–2186. doi:10.3390/ijms19082166

36. Mondal S, Mason RE, Huggins T, Hays DB. QTL on wheat (*Triticum aestivum* L.) chromosomes 1B, 3D and 5A are associated with constitutive production of leaf cuticular wax and may contribute to lower leaf temperatures under heat stress. Euphytica. 2015; 201: 123–130. doi:10.1007/s10681-014-1193-2

37. Jha UC, Bohra A, Singh NP. Heat stress in crop plants: Its nature, impacts and integrated breeding strategies to improve heat tolerance. Plant Breed. 2014; 133: 679–701.

38. Wang YH, Song H, Quan QS, Wang YE, Zhang XQ, Yan LY, et al. A method of heat tolerance identification and selection of germplasm in bottle gourd type rootstocks. Chinese Patent No 2012100939951. 2014; Beijing: National Intellectual Property Administration, PRC.

39. Zhou WJ, Leul M. Uniconazole-induced alleviation of freezing injury in relation to changes in hormonal balance, enzyme activities and lipid peroxidation in winter rape. Plant Growth Regul. 1998; 26: 41–47.

40. He AL, Niu SQ, Zhao Q, Li YS, Gou JY, Gao HJ, et al. Induced salt tolerance of *Perennial ryegrass* by a novel bacterium strain from the rhizosphere of a desert shrub *Haloxylon ammodendron*. Int J Mol Sci. 2018; 19: 469–472. doi:10.3390/ijms19020469

41. Sun CX, Cao HX, Chen ST, Feng ML, Li RS, Ma ZL. Study on cold resistance of snake fruit by application of electrical conductivity and logistic equation. Acta Agr Jiangxi. 2009; 21: 33–35 (in Chinese).

42. Ling F, Li C, Hui Z, Jiao J, Lyu P. Measurement of cold tolerance by electrical conductivity method in associated with the logistic equation on different varieties of *Oleaeuropaea* L. Acta Agr Guangdong. 2015; 1: 13–17 (in Chinese). doi:10.19386/j.cnki.jxnyxb.2009.04.010

43. Saadalla MM, Shanahan JF, Quick JS. Heat tolerance in winter wheat: I. hardening and genetic effects on membrane thermostability. Crop Sci. 1990; 30: 1243–1247. doi: 10.2135/cropsci1990.0011183X003000060017x

44. Sui Y, Miao Y, Hu N, Zhao Y, Zhou Y. Genetic analyzing yield traits of chilli pepper. J Nanjing Agri Univ. 2014; 37: 46–52. doi:10.7685/j. issn. 1000-2030.2014.04.007

45. Murray M, Thompson WF. Rapid isolation of high molecular weight plant DNA. Nucleic Acids Res. 1980; 8: 4321–4326.

46. Soldi S, Vasileiadis S, Uggeri F, Campanale M, Morelli L, Fogli MV, et al. Modulation of the gut microbiota composition by rifaximin in non-constipated irritable bowel syndrome patients: a molecular approach. Clin Exp Gastroenterol. 2015; 8: 309–325.doi: 10.2147/CEG.S89999

47. Li H, Durbin R. Fast and accurate short read alignment with Burrows-Wheeler transform. Bioinformatics. 2009; 25: 1754–1760.

48. Zuryn S, Gras LS, Jamet K, Jarriault S. A strategy for direct mapping and identification of mutations by whole-genome sequencing. Genetics. 2010; 186: 427–430. doi: 10.1534/genetics.110.119230

49. Abe A, Kosugi S, Yoshida K, Natsume S, Takagi H, Kanzaki H, et al. Genome sequencing reveals aronomically important loci in rice using MutMap. Nature Biotech. 2012; 30: 174–178.

50. Pruitt KD, Tatusova T, Maglott DR. NCBI reference sequences (RefSeq): a curated non-redundant sequence database of genomes, transcripts and proteins. Nucleic Acids Research. 2007; 35: 61–65.

51. Gull IS, Hulpiau P, Saeys Y, Roy F. Metazoan evolution of the armadillo repeat super family. Cell Mol Life Sci. 2017; 74: 525–541.

52. Lichocka M, Rymaszewski W, Morgiewicz K, Barymow-Filoniuk I, Chlebowski A, Sobczak M, et al. Nucleus- and plastid-targeted annexin 5 promotes reproductive development in *Arabidopsis* and is essential for pollen and embryo formation. BMC Plant Biol. 2018; 18: 183–198. doi: 10.1186/s12870-018-1405-3

53. Hruba P, Honys D, Twell D, Capkova V, Tupy J. Expression of β-galactosidase and β-xylosidase genes during microspore and pollen development. Planta. 2005; 220: 931–940.

54. Wang CG, Chen XQ, Lan TY, Li H, Song WQ. Cloning and transcript analyses of the chimeric gene associated with cytoplasmic male sterility in cauliflower (*Brassica oleracea* var. *botrytis*). Euphytica. 2006; 151: 111–119.

55. Idänheimo N, Gauthier A, Salojärvi J, Siligato R, Brosché M, Kollist H, et al. The Arabidopsis thaliana cysteine-rich receptor-like kinases CRK6 and CRK7 protect against a poplastic oxidative stress. Biochem Bioph Res Co. 2014; 445: 457–462.

56. Umate P, Tuteja R, Tuteja N. Genome-wide analysis of helicase gene family from rice and *Arabidopsis*: a comparison with yeast and human. Plant Mol Biol. 2010; 73: 449–465.

57. Xiao WF, Chen P, Xiao JS, Wang L, Liu TH, Wu YF, et al. Comparative transcriptome profiling of a thermal resistant vs. sensitive silkworm strain in response to high temperature under stressful humidity condition. PLoS One. 2017; 12: e0177641. doi: org/10.1371/journal.pone.0177641

58. Yabuta S, Ifuku K, Takabayashi A, Ishihara S, Ido K, Ishikawa N, et al. Three PsbQ-Like proteins are required for the function of the chloroplast NAD (P) H dehydrogenase complex in *Arabidopsis*. Plant Cell Physiol. 2010; 51: 866–876.

59. Huang N, Ling H, Su YC, Liu F, Xu LP, Su WH, et al. Transcriptional analysis identifies major pathways as response components to Sporisorium scitamineum stress in sugarcane. Gene. 2018; 678: 207–218.

60. Wang Y, Cai SY, Yin LL, Shi K, Xia XJ, Zhou YH, et al. Tomato HsfA1a plays a critical role in plant drought tolerance by activating ATG genes and inducing autophagy. Autophagy. 2015; 11: 2033–2047.

61. Rashkova S, Karam SE, Pardue ML. Element-specific localization of drosophila retrotransposon gag proteins occurs in both nucleus and cytoplasm. Proc Natl Acad Sci USA. 2002; 99: 3621–3626.

62. Donlin MJ, Lisch D, Freeling M. Tissue-specific accumulation of MURB, a protein encoded by MuDR, the autonomous regulator of the mutator transposable element family. Plant Cell. 1995; 7: 1989–2000.

63. Abeliovich H, Tzfati Y, Shlomai J. A trypanosomal CCHC-Type zinc finger protein which binds the conserved universal sequence of kinetoplast DNA minicircles: isolation and analysis of the complete cDNA from *Crithidia fasciculata*. Mol Cell Biol. 1993; 13: 7766–7773.

64. Kraus B, Boller K, Reuter A, Schnierle BS. Characterization of the human endogenous retrovirus K Gag protein: identification of protease cleavage sites. Retrovirology. 2011; 8: 21–29.

65. Cowles CR, Odorizzi G, Payne GS, Emr SD. The AP-3 adaptor complex is essential for cargo-selective transport to the yeast vacuole. Cell. 1997; 91: 109–118.

66. Parsiegla G, Noguere C, Santell L, Lazarus RA, Bourne Y. The structure of human DNase I bound to magnesium and phosphate ions points to a catalytic mechanism common to members of the DNase I-like super family. Biochemistry. 2012; 51: 10250–10258.

67. Du YL, Singh R, Alkhalaf LM, Kuatsjah E, He HY, Eltis LD, Ryan KS. A pyridoxal phosphate-dependent enzyme that oxidizes an unactivated carbon-carbon bond. Nat Chem Biol. 2016; 12: 194–199.

68. Kaya H, Ichi-Shibahara K, Taoka KI, Iwabuchi M, Stillman B, Araki T. FASCIATA genes for chromatin assemblyfactor-1 in *Arabidopsis* maintain the cellular organization of apical meristems. Cell. 2001; 104: 131–142.

69. Hirayama T, Shinozaki K. Research on plant abiotic stress responses in the post-genome era: past, present and future. Plant J. 2010; 61: 1041–1052. doi:10.1111/j.1365-313X.2010.04124.x

70. Paliwal R, Reoder MS, Kumar U, Srivastava JP, Joshi AK. QTL mapping of terminal heat tolerance in hexaploid wheat (*T*. *aestivum* L.). Theor Appl Genet. 2012; 125: 561–575. doi:10.1007/s00122-012-1853-3

71. Barakat MN, Al-Doss AA, Elshafei AA, Moustafa KA. Identification of new microsatellite marker linked to the grain filling rate as indicator for heat tolerance genes in F2 wheat population. Aust J Crop Sci. 2011; 5: 104–110.

72. Talukder SK, Md A, Babar K, Vijayalakshmi J, Poland PVV, Bowden R, et al. Mapping QTL for the traits associated with heat tolerance in wheat (*Triticum aestivum* L.). BMC Genet. 2014; 15: 97–110. doi:10.1186/s12863-014-0097-4

73. Farnham M, Björkman T. Evaluation of experimental broccolihybrids developed for summer production in the Eastern United States. Hort Science. 2011; 46: 858–863.

74. Branham SE, Stansell ZJ, Couillard DM, Farnham MW. Quantitative trait loci mapping of heat tolerance in broccoli (*Brassica oleracea* var. *italica*) using genotyping-by-sequencing. Theor Appl Genet. 2017; 130: 529–538. doi: 10.1007/s00122-016-2832-x

75. Bouwkamp JC, Summers WL. Inheritance of resistance to temperature-drought stress in the snap bean. J Hered. 1982; 73: 385–386. doi:10.1093/oxfordjournals.jhered.a109680

76. Marfo KO, Hal AE. Inheritance of heat tolerance during pod set in cowpea. Crop Sci. 1992; 32: 912–918. doi: 10.2135/cropsci1992.0011183X0032000400115x

77. Li XM, Chao DY, Wu Y, Huang X, Chen K, Cui LG, et al. Natural alleles of a proteasome a2 subunit gene contribute to thermo tolerance and adaptation of African rice. Nat Genet. 2015; 47: 827–833. doi:10.1038/ng.3305

78. Singh D, Singh CK, Tomar RSS, Pal M. Genetics and molecular mapping of heat tolerance for seedling survival and pod set in lentil. Crop Sci. 2017; 57: 3059–3067. doi: 10.2135/cropsci2017.05.0284

79. Rahamana M, Mamidib S, Rahmana M. Genome-wide association study of heat stress tolerance traits in spring-type *Brassica napus* L. under controlled conditions. Crop J. 2018; 6: 115–125.

80. Wahid A, Gelani S, Ashraf M, Foolad MR. Heat tolerance in plants: An overview. Environ Exp Bot. 2007; 61: 199–223.

81. Hamada A, Zintal G, Hegab MM, Pandey R, Asard H, Abuelsoud W. High salinity induces different oxidative stress and antioxidant responses in maize seedlings organs. Front Plant Sci. 2016; 26: 1–11. doi: 10.3389/fpls.2016.00276

82. Ye CR, Tenoriol FA, Argayoso MA, Laza MA, Koh HJ, Redoña ED, et al. Identifying and confirming quantitative trait loci associated with heat tolerance at flowering stage in different rice populations. BMC Genet. 2015; 16: 41–51. doi: 10.1186/s12863-015-0199-7

83. Davies JM. Annexin-mediated calcium signalling in plants. Plants (Basel). 2014; 3: 128–140.

84. Laohavisit A, Davies JM. Annexins. New Phytol. 2011; 189: 40–53.

85. Laohavisit A, Brown AT, Cicuta P, Davies JM. Annexins: components of the calcium and reactive oxygen signaling network. Plant Physiol. 2010; 152: 1824–1289.

86. Zhu J, Wu X, Yuan S, Qian D, Nan Q, An L, et al. Annexin5 plays a vital role in *Arabidopsis* pollen development via Ca2+-dependent membrane trafficking. PLoS One. 2014; 9: e102407.

87. Marks MS, Ohno H, Kirchhausen T, Bonifacino JS. Protein sorting by tyrosine-based signals. Trends Cell Biol. 1997; 7: 124–128.

88. Fan L, Hao H, Xue Y, Zhang L, Song K, Ding Z, Botella MA, Wang H, Lin J. Dynamic analysis of *Arabidopsis* AP2 s subunit reveals a key role in clathrin-mediated endocytosis and plant development. Development. 2013; 140: 3826–3837.

89. Park M, Song K, Reichardt I, Kim H, Mayer U, Stierhof YD, et al. Arabidopsis u-adaptin subunit AP1M of adaptor protein complex 1 mediates late secretory and vacuolar traffic and is required for growth. Proc Natl Acad Sci USA. 2013; 110: 10318–10323.

90. Bassham DC, Brandizzi F, Otegui MS, Sanderfoot AA. The secretory system of *Arabidopsis*. Arabidopsis Book. 2008; 6: e0116.

91. Simpson F, Bright NA, West MA, Newman LS, Darnell RB, Robinson MS. A novel adaptor-related protein complex. J Cell Biol. 1996; 133: 749–760.

92. Feraru E, Paciorek T, Feraru MI, Zwiewka M, De Groodt R, De Rycke R, et al. The AP-3 β adaptin mediates the biogenesis and function of lytic vacuoles in *Arabidopsis*. Plant Cell. 2010; 22: 2812–2824.

93. Zwiewka M, Feraru E, Möller B, Hwang I, Feraru MI, Kleine-Vehn J, et al. The AP-3 adaptor complex is required for vacuolar function in *Arabidopsis*. Cell Res. 2011; 21: 1711–1722.

94. Feng QN, Liang X, Li S, Zhang Y. The adaptor protein-3 complex mediates pollen tube growth by coordinating vacuolar targeting and organization. Plant Physiol. 2018; 177: 216–225.

95. Holzman LB, Merritt SE, Fan G. Identification, molecular cloning, and characterization of dual leucine zipper bearing kinase. J Biol Chem. 1994; 269: 30808–30817.

96. Umezawa T, Yoshida R, Maruyama K, Yamaguchi-Shinozaki K, Shinozaki K. SRK2C, a SNF1-related protein kinase 2, improves drought tolerance by controlling stress-responsive gene expression in *Arabidopsis thaliana*. Proc Natl Acad Sci USA. 2004; 101: 17306–17311. doi: 10.1073pnas.0407758101

